# Dysregulation of Pulmonary Responses in Severe COVID-19

**DOI:** 10.1101/2020.10.23.353177

**Authors:** Dandan Wu, Xuexian O. Yang

**Affiliations:** Department of Molecular Genetics and Microbiology, University of New Mexico School of Medicine, Albuquerque, NM 87131, USA; College of Agronomy, Hunan Agricultural University, Changsha, Hunan 410128, China

**Keywords:** COVID-19, severity, TH17, IgA, type I interferon, PAI-1

## Abstract

Patients with coronavirus disease 2019 (COVID-19) predominantly have a respiratory tract infection with various symptoms and high mortality is associated with respiratory failure second to severe disease. The risk factors leading to severe disease remain unclear. Here, we reanalyzed a published single-cell RNA-Seq (scRNA-Seq) dataset and found that bronchoalveolar lavage fluid (BALF) of patients with severe disease compared to those with mild disease contained decreased TH17 type cells, decreased IFNA1 expressing cells with lower expression of toll-like receptor 7 (TLR7) and TLR8, increased IgA expressing B cells, and increased hyperactive epithelial cells (and/or macrophages) expressing matrix metalloproteinases (MMPs), Hyaluronan synthase 2 (HAS2), and Plasminogen activator inhibitor-1 (PAI-1), which may together contribute to the pulmonary pathology in severe COVID-19. We propose IFN-I (and TLR7/TLR8) and PAI-1 as potential biomarkers to predict the susceptibility to severe COVID19.

## Introduction

COVID-19, a SARS-CoV-2 caused infectious disease, manifests various symptoms ranging from asymptomatic, mild to very severe and leads to multiple organ injury and even death. Poor outcomes are associated with older age (especially over 65) and underlying conditions including diabetes, cardiovascular disease, hypertension, obesity, and chronic obstructive pulmonary disease (COPD) (1). Heightened serum levels of IL-6, CRP and D-dimer, lymphopenia, neutrophilia, and other complications have been reported in sever COVID-19 (2,3). In severe cases, cytokine release syndrome (also called “cytokine storm”) results in acute respiratory distress syndrome (ARDS) with drowning edema in the lung, which is broadly accepted as one of the major causes of death. Recent studies showed that pre-existing, cross-reactive T cells (elicited by prior infection with “common cold” coronaviruses) might limit the disease severity (4–6). In addition to T cells, cross-reactive antibodies are also present in unexposed healthy cohorts, especially those aged 6-16 years (7). However, the pre-existing immunity cannot explain the susceptibility in people aged 65 and above because they have more chances to be infected with “common cold” coronaviruses than young people and may have persistent cross-reactive T cells. It cannot explain that men are more susceptible than women, either. Identifying risk factors that drive the transition to severe disease remains highly demanded and would benefit the treatment, prevention, and vaccination. Here, we reanalyzed a published scRNA-Seq dataset and found that bronchoalveolar lavage fluid (BALF) of patients with severe disease compared to those with mild disease exhibited dysregulation of TH17 type cells, IgA expressing B cells, type I interferon (IFN-I) pathway, and matrix metalloproteinases (MMPs), Hyaluronan synthase 2 (HAS2), and Plasminogen activator inhibitor-1 (PAI-1) expressing cells.

## Main

### Severe COVID-19 displays decreased TH17 type cells and increased IgA^+^ B in BALFs

Peripheral blood mononuclear cell (PBMC) studies have demonstrated dysregulated myeloid (monocyte and neutrophil) and CD8^+^ T cell compartments in severe COIVD-19 (3,8,9). Currently, there are only a few studies focusing on lung local responses. Zhou et al. revealed a hyper-proinflammatory gene expression profile by meta-transcriptomic sequencing of BALF cells (10). Compared to community-acquired pneumonia patients and healthy controls, BALF cells of COVID-19 patients highly express proinflammatory genes, especially chemokines, suggesting that SARS-CoV-2 infection causes hypercytokinemia. Like SARS-CoV, SARS-CoV-2 robustly triggered expression of numerous IFN-stimulated genes (ISGs). Liao et al. compared BALF cell responses in mild and severe COVID-19 cases using scRNA-Seq (11). The BALFs of severe cases had more abundant macrophages and neutrophils with a decrease in the CD8^+^ T cell population, and expressed elevated levels of cytokines, IL1B, IL6, and TNF, and chemonkines, compared with those of the mild cases. By leveraging Liao et al.’s scRNA-Seq dataset, we further evidenced the dysregulation of T helper (TH) cells, B cells, IFN-I pathway, and tissue factors in the severe cases.

The scRNA-Seq dataset (GEO accession number GSE145926) (11), including 3 healthy controls, 3 mild cases, and 6 severe cases, was downloaded and analyzed using SeqGeq software (FlowJo LLC) and one-side unpaired T-test was used to calculate the statistical significance. The focus was on the comparison between mild and severe cases; healthy controls were included as references. In the CD4+ TH cell compartment, there were no significant differences in TH1 (TBX21^+^), TH2 (GATA3^+^), and regulatory T (FOXP3^+^) cells between the mild vs. severe cases (Fig. 1A). Interestingly, compared with mild cases, BALFs of severe cases had decreased RORC^+^ or CCR6^+^ TH17 cells (Fig. 1A) and γδT (TRDC^+^) cells (Fig. 1B); the latter also express TH17 type cytokines, IL17 and IL17F (and TH1 type cytokine IFNγ). Although TH17 cells are considered as a potent mediator of tissue pathology, they are essential in anti-viral immunity through promoting TH1, cytotoxic T lymphocyte, B cell responses and are implicated in combating concomitant bacterial (and maybe also fungal) infection (12,13). The impaired TH17 responses in severe cases suggest a protective role of TH17 type cells, which further implicate potential benefit of antibiotics (and maybe also antimycotics) for patients with severe disease. Besides the lung, the intestine, another major mucosal site, has active TH17 responses. SARS-CoV-2 also infects the intestine, where express its receptors ACE2 and TMPRSS2 (14). A large number of CD4^+^ CCR6^+^ TH17 cells have been reported in PBMCs of a deceased patient (15). In addition, there are more SARS-CoV-2 reactive TH17 cells highly expressing IL17 (IL17A) and CCR6 in PBMCs of hospitalized than non-hospitalized patients (16). Therefore, the systemic role of TH17 cells in the disease progress, especially the development of ARDS, need further define. Interestingly, 4 out of 6 BALF samples of severe cases expressed IL22 whereas none of mild cases expressed detectable levels of IL22 (Fig. 1C). IL22^+^ cells were CD3E^+^ CD4^+^ AHR^+^ but TRDC^−^ and RORC^−^, therefore belonging to TH22 but not TH17 cells. Whether IL22 plays a role in the disease severity remains to be determined. Besides the dysregulation in the T cell compartment, severe cases had increased frequencies of IgA1 (IGHA1^+^) [and a trend of increasing IgG1 (IGHG1 ^+^)] expressing B cells (Fig. 1D). Generally, antibodies confer favorable humoral immunity; however, massive immune complexes can be a driving force of tissue permeability as seen in systemic lupus erythematosus, in agreement with Chen et al.’s observation that higher virus-specific antibody titers correlate with disease severity (17). In summary, decreased TH17 type T cells and increased IgA secreting B cells may augment the disease severity.

**Figure 1.**
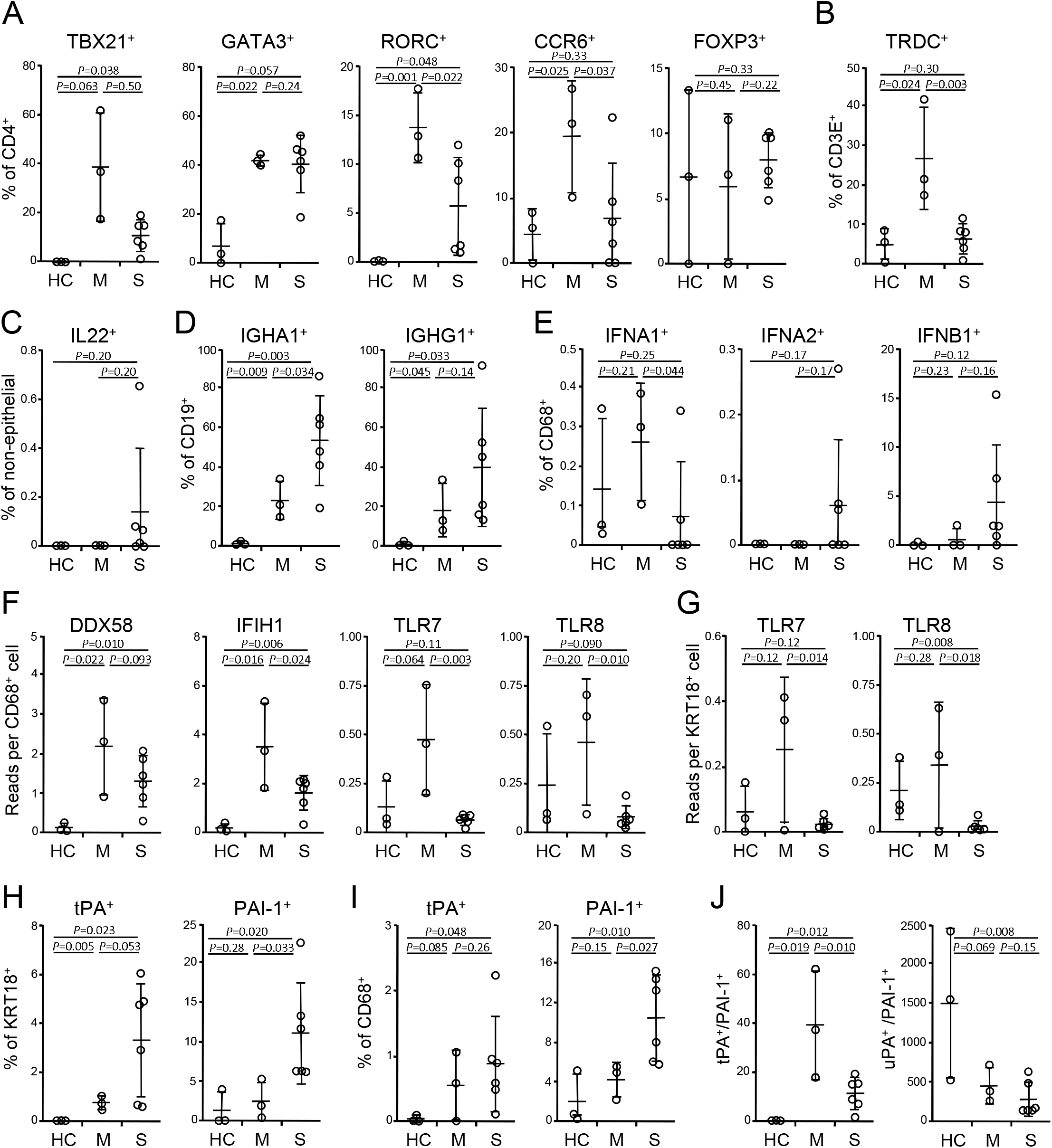
Dysregulation of TH and B cells, IFN-I, and tissue factors in BALFs. (A) Frequencies of TH1 (TBX21^+^), TH2 (GATA3^+^), TH17 (RORC^+^ or CCR6^+^) cells and regulatory T cells (FOXP3^+^) in BALF cells on a CD4^+^ CD14^−^ gate. (B) Frequencies of γδT cells (TRDC^+^) on a CD3E^+^ gate. (C) Frequencies of IL22^+^ cells on a KRT18^−^ gate. (D) Frequencies of IgA1 (IGHA1) and IgG1 (IGHG1) expressing B cells on a CD19^+^ gate. (E) Frequencies of IFN-I expressing macrophages on a CD68^+^ gate. (F, G) Abundances of RNA recognition receptors in macrophages (F) and epithelial cells (G). (H, I) Frequencies of tPA and PAI-1 expressing epithelial cells (H) and macrophages (I). (J) Ratios of total tPA vs PAI-1 and total uPA vs PAI-1 expressing cells. HC, healthy control (n = 3); M, mild (n = 3); S, severe (n = 6). Mean and s.d. were shown. *p* values, unpaired T-test.

### Severe COVID-19 has an impaired IFN-I response

IFN-Is play an important role in anti-viral immunity. Due to sequencing depth, IFN-Is were detected mainly in CD68^+^ macrophages and a few other cells, such as KRT18^+^ epithelial cells. Macrophages of severe cases had decreased frequencies of IFNA1 but not IFNA2 and IFNB1 expressing cells relative to those of mild cases (Fig. 1E). Consistently, patients with severe disease had a decrease of expression of RNA sensors, TLR7 and TLR8 in both macrophages and epithelial cells and IFIH1 (encoding MDA5) but not DDX58 (encoding RIG-I) in macrophages (Fig. 1F, G). There were no differences in IFIH1 and DDX58 in epithelial cells in both groups (data not shown). These results implicate an essential role of IFN-I pathway in the disease susceptibility, in agreement with previous observations (18,19). Although IFN-I only marginally decline with aging and this decline can recover around age of 55 (20), during viruses challenge (such as influenza virus and West Nile virus), both pDCs and mDCs of aged donors have a decreased capacity of IFN-I production, which subsequently impairs the secretion of IFNγ in CD4 T cells and IFNγ, perforin and granzyme in CD8 T cells (21–23). In addition, certain diseases, such as diabetes and hypertension, can also impair IFN-I production (24). Furthermore, a recent study showed that women had higher IFNA2 levels than men (25). In summary, the defect of IFN-I pathway, associated with advancing age, gender, and some diseases, emerges as a risk factor of severe COVID-19.

### Severe COVID-19 exhibits enhanced expression of MMPs

MMPs degrade extracellular matrix components of the intersititium and tight junction and therefore, increase alveolar permeability that is observed in many destructive lung diseases, including ARDS, COPD, tuberculosis, sarcoidosis and IPF, whereas tissue inhibitors of metalloproteinases (TIMPs) inhibit the enzymatic activity of MMPs (26,27). Of abundant MMPs and TIMPs, BALF epithelial cells of severe COVID-19 had elevated frequencies of MMP7^+^ [mild vs. severe (mean ± s.d.), 2.94 ± 1.24% vs. 13.19 ± 8.48%, *p* = 0.042] and MMP9^+^ (0.25 ± 0.43% vs. 1.60 ± 1.18%, *p* = 0.053) cells and a trend of increasing MMP2^+^ (0% vs. 1.02 ± 0.98%, *p* = 0.062) and MMP13^+^ (0.72 ± 0.30% vs. 5.39 ± 5.15%, *p* = 0.084) portions compared with mild cases, but no differences were found in MMP14^+^ (27.44 ± 13.86% vs. 24.49 ± 11.24%, *p* = 0.37) cell percentages. There were no alterations in the frequencies of either TIMP1^+^ (27.44 ± 13.86% vs. 24.49 ± 11.24%, *p* = 0.37) or TIMP2^+^ (26.27 ± 20.24% vs. 24.49 ± 11.24%, *p* = 0.43) expressing epithelial cells between mild vs. severe cases. In addition to epithelial cells, macrophages also expressed MMPs and the expression patterns were similar with those of epithelial cells but only MMP9 (1.01 ± 0.63% vs. 5.03 ± 3.54%, *p* = 0.053) reached significance. Macrophages of severe cases had higher percentages of TIMP1^+^ cells (64.47 ± 13.03% vs. 77.42 ± 15.18%, *p* = 0.040) whereas a trend in decreasing TIMP2^+^ frequencies (70.61 ± 17.99% vs. 40.30 ± 8.77%, *p* = 0.098) relative to those of mild cases. Together, over-expression of MMPs, in addition to macrophage and neutrophil products, such as ROS, NO, and enzymes, may promote intensive lung tissue destruction in severe COVID-19.

### Severe COVID-19 manifests increased expression of Mucin 1 (MUC1), HAS2 and PAI-1

Hypersecretion of MUCs is a common feature of many lung diseases, including viral infection, asthma, and COPD. Of abundant mucins, MUC1 (1.39 ± 0.59 vs. 5.02 ± 4.05 reads/cell, *p* = 0.090) expression is upregulated (but did not reach significance) in epithelial cells of severe vs. mild COVID-19, whereas there were no differences in MUC4 (4.57 ± 2.44 vs. 5.49 ± 3.81 reads/cell, *p* = 0.36) and MUC16 (8.34 ± 5.17 vs. 5.22 ± 3.07 reads/cell, *p* = 0.14) between the two groups. Hypersecretion of hyaluronic acid presents in severe COVID-19 (28,29). Consistently, HAS2 (0.08 ± 0.10% vs. 0.39 ± 0.27%, *p* = 0.042) but not HAS3 (1.21 ± 0.69% vs. 3.01 ± 2.75%, *p* = 0.16) was upregulated in epithelial cells of severe cases, whereas HAS1 expressing cells were very few (data not shown). In addition to hyper-production of MUC1 and hyaluronic acid, there were more epithelial cells producing tPA and PAI-1 in severe disease compared with mild disease (Fig. 1H). Severe cases also had higher frequencies of PAI-1^+^ but not tPA^+^ macrophages (Fig. 1I). Total tPA^+^ cells to PAI-1^+^ (tPA/PAI-1) cell ratios were decreased in severe relative to mild cases and there was no difference in uPA/PAI-1 (Fig. 1J). tPA and uPA promote fibrinolysis, whereas PAI-1 inhibits this process. Decreased tPA/PAI-1 in severe cases suggests that dysregulation of coagulation plays an important role in the disease severity, in agreement with Tang et al.’s study that abnormal coagulation leads to poor prognosis COVID-19 (30). Heightened expression of tPA and PAI-1 was also reported in highly lethal acute hantavirus cardiopulmonary syndrome (31). PAI-1 levels are inversely correlated with D-dimer concentrations. PAI-1 expression is induced by a number of proinflammatory cytokines, such as IL1, IL6, TNF, and TGFβ, and hormones, such as insulin, glucocorticoids, and adrenaline (reviewed in refs. (32,33)). Advancing age is considered as a major contributor to increased expression of PAI-I (34), which is correlated with aging-associated cardiovascular diseases and metabolic syndromes, such as atherosclerosis, hypertension, obesity, and diabetes (32,33). Moreover, men have higher serum PAI-1 levels than women (35,36). Therefore, elevation of PAI-I expression can serve as a risk factor of severe COVID-19, especially in people with older age, male gender, and/or a group of preexisting conditions. In summary, augmented expression of MUC1, HAS2 and PAI-1 is associated with more severe disease and may contribute to the drowning edema and the coagulation disorder in severe COVID-19.

## Conclusion

Dysregulation of pulmonary responses, including decreased TH17 cells (and CD8^+^ T cells (11)) and IFN-I (the latter is associated with impaired TLR7 and TLR8 expression), and elevation of IgA, MMPs, MUC1, hyaluronic acid, and PAI-I (as well as myeloid cells (11)), is correlated with the disease severity (see more discussion in Fig. 2). Amongst above mentioned factors, IFN-I and PAI-1 are dysregulated in older age, male gender, and preexisting diseases that are associated with increased risk to develop severe disease, we propose IFN-I (and TLR7/TLR8) and PAI-1 as biomarkers to predict the susceptibility to severe COVID19 (and maybe also other lung infections), even in uninfected populations.

**Figure 2.**
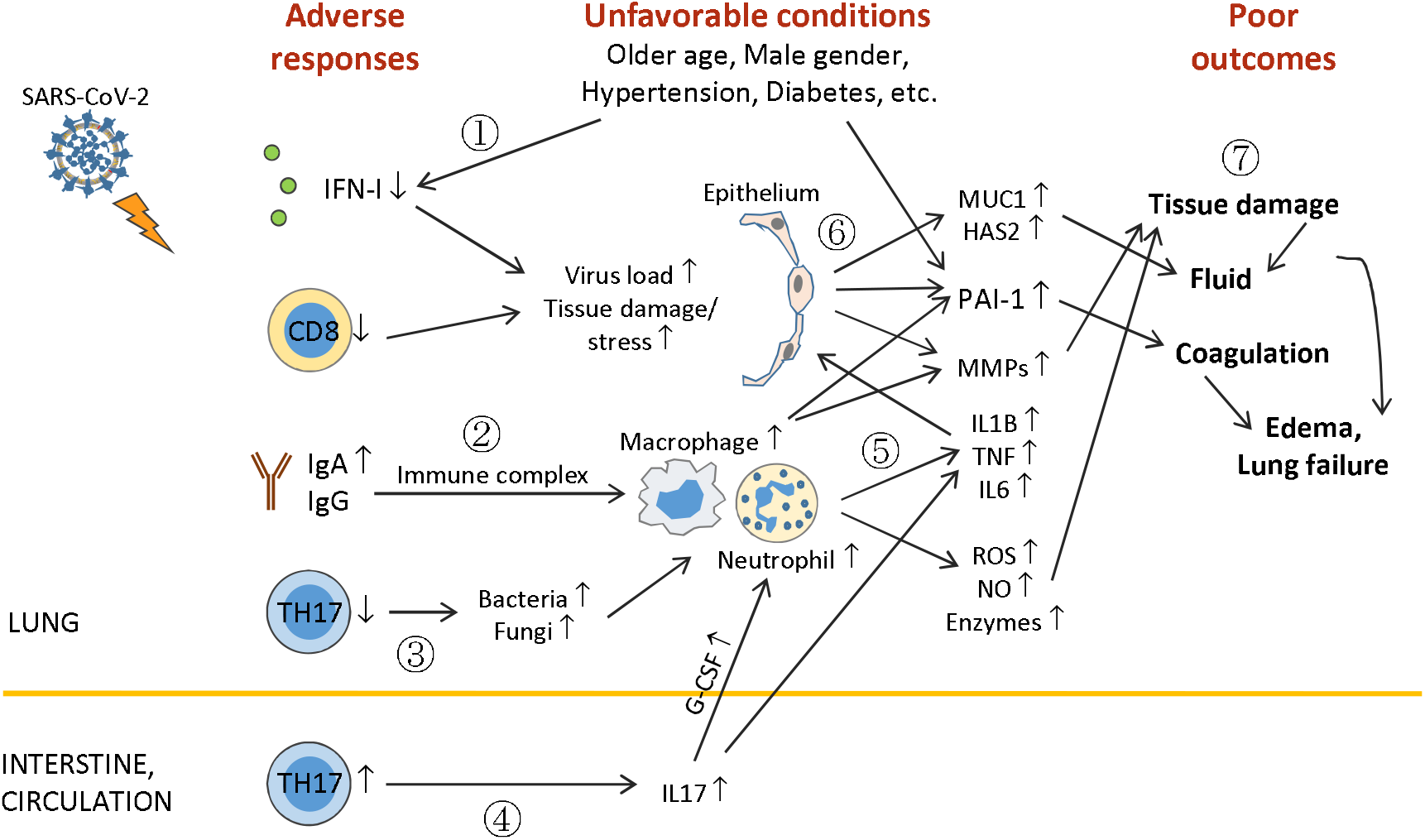
Outline of unfavorable conditions and deleterious pulmonary responses. (1) Older age, male gender, underlying conditions (such as hypertension, diabetes, etc.), and unknown factors (including genetic background) impair anti-viral immunity (including IFN-I deficiency and decreased CD8^+^ T and TH17 cells), leading to higher virus loads and tissue damage/stress. (2) Elevation of humoral responses results in massive immune complexes that activates macrophages and neutrophils. (3) Decreased TH17 cell responses cause overgrowth of commensal bacteria and fungi, which further activate macrophages and neutrophils. (4) TH17 hyper-activation and/or expansion in the intestine (?) cause high levels of serum IL17, which induces G-CSF expression and in turn, promotes neutrophilia. (5) Hyper-activated macrophages and neutrophils release immense amounts of proinflammatory cytokines, leading to cytokine release syndrome and subsequent ARDS, and tissue destructive products, such as ROS, NO, MMPs, and other enzymes. (6) During ARDS, proinflammatory cytokines act on epithelial cells and induce MMPs, mucins, hyaluronic acids, antimicrobial peptides, and PAI-1 (unfavorable conditions also elevate PAI-1 expression). (7) ROS, NO, MMPs, and other enzymes cause epithelial and endothelial leakage, leading to tissue fluid/plasma accumulation in alveolar spaces. Mucins, hyaluronic acids, and antimicrobial peptides concentrate alveolar fluids and thicken mucosal lining, resulting in drowning edema and even lung failure. Heightened PAI-1 facilitates coagulation and strengthens edema formation (and thrombosis). ARDS also has systemic consequences causing multi-organ damage.

## Acknowledgments

XOY was supported by NIH AI142200 and HL148337.

